# Oncogenic metabolic rewiring independent of proliferative control in human mammary epithelial cells

**DOI:** 10.1101/2022.04.08.486845

**Authors:** Wentao Dong, Mark A. Keibler, Sun Jin Moon, Patricia Cho, Nian Liu, Christian J. Berrios, Joanne K. Kelleher, Hadley D. Sikes, Othon Iliopoulos, Jonathan L. Coloff, Matthew G. Vander Heiden, Gregory Stephanopoulos

## Abstract

The use of isotopic tracers and metabolic flux analysis (MFA) has unveiled a number of metabolic pathways differentially activated in cancer cells. To support efforts to design effective metabolic therapies for cancer, we sought to distinguish metabolic behavior in cancer versus normal cells growing at the same rate. To this end, we performed ^13^C-isotope tracing and MFA in human mammary epithelial cells (HMECs) harboring different combinations of oncogenes. By introducing a new quantity termed metabolic flux intensity, defined as pathway flux divided by specific growth rate, we showed that metabolism is dually controlled by proliferation and oncogenotypes. ^13^C-MFA further revealed that oxidative pentose phosphate pathway (oxPPP), malate dehydrogenase (MDH) and isocitrate dehydrogenase (IDH) were most enhanced in cancerous HMECs. Drug targeting of these pathways selectively reduced growth in the tumorigenic HMEC line. Our study provides direct evidence that metabolism of cancer cells is different than that of normal proliferating cells.

## Introduction

Metabolic rewiring is one of the hallmarks of cancer ^1^. Our knowledge of such changes has now extended beyond the Warburg effect ^2–8^, which refers to the paradoxical phenomenon that cancer cells preferentially metabolize glucose to lactate regardless of oxygen availability ^9^. Over the past decades, various types of metabolic alterations in cancer cells have been documented ^7, 10–15^, prompting extensive investigation of the link between reprogrammed cell signaling pathways and rewired cellular metabolism ^16–21^. In addition, drug targeting of certain metabolic pathways has been demonstrated to be a promising cancer treatment strategy ^22–24^. Despite this progress, a fundamental question remains unanswered: whether there is a difference in the metabolism of cancer cells and their normal proliferating counterparts. This deficiency has hindered our understanding of cancer metabolism and the efforts to develop effective cancer therapies targeting metabolism with reduced side effects.

This question has not been directly addressed before. It has been speculated that rewired metabolic behavior associated with cancer may be a simple manifestation of reprogrammed cellular energetics required to accommodate deregulated cell growth ^1, 25, 26^. For example, the Warburg effect is observed in both tumor and highly proliferative normal cells ^1, 27, 28^. Several types of proliferating immune cells and stem cells rely on the glycolytic phenotype, a metabolic hallmark of cancer, to maintain their respective functionality and lineage ^29–33^. Furthermore, both cancer and normal proliferating cells may selectively express the M2 isoform of pyruvate kinase, which is capable of promoting anabolism and cellular growth ^10, 34^. Reductive carboxylation of glutamine has also been reported in both proliferating endothelial cells ^35^ and several types of cancer cells ^36, 37^, suggesting that rewired glutamine metabolism is not unique to cancer.

However, the notion that metabolic behavior in cancer and normal proliferative cells is equivalent is still questionable. For example, comparison between normal proliferative and tumorigenic liver cells revealed that the latter exhibited upregulated glutaminase and transaminase activities ^24^. In addition, copy number amplification of phosphoglycerate dehydrogenase was reported in melanoma relative to normal tissue counterparts ^38, 39^. Moreover, increased expression of proline oxidase has been observed in pancreatic ductal adenocarcinoma compared to normal pancreatic cells ^40^. Although these findings suggest that cancer metabolism may not be simply regarded as rewired proliferative energetics, the evidence is still inconclusive: the proliferation rate of cancer cells in these studies was different than that of the normal cells. Therefore, the metabolic alteration due to proliferation introduces a confounding factor in the observed metabolic shifts. This challenge limits our ability to conclusively determine whether metabolism of cancer cells is truly different than that of normal proliferating cells.

In order to address this question, we developed a panel of HMECs from the same genetic background yet harboring different combinations of oncogenic manipulations. Through the use of ^13^C-MFA and the introduction of a new quantity referred to as metabolic flux intensity (MFI), we decoupled proliferative control of metabolism and removed cellular growth as a factor affecting metabolic rewiring. In addition, ^13^C-isotopic labeling analysis and MFA also revealed a distinct metabolic pattern within the tricarboxylic acid (TCA) cycle and lipogenesis. Our results suggest that metabolism is directly controlled by both oncogenotypes and proliferation. The proliferation-independent metabolic shifts identified in our work support the effort to design cancer therapies that target specific metabolic pathways related with tumors only and not affecting growth of healthy cells.

## Results

### Construction of HMECs with different combinations of oncogenes

To compare cancer and proliferative metabolism, we developed a panel of HMECs harboring different oncogenes. The resulting HMECs share the same genetic background from the parental cell: an immortalized, yet non-cancerous, HMEC 184A1 (ATCC), the growth of which is dependent on the amount of epidermal growth factor (EGF) present in media. We then overexpressed defined oncogenes and dominant negative mutant forms of tumor suppressors in the original HMEC 184A1.

Based on the characteristic properties of cancer cells ^41^, we considered three key attributes relevant to our specific *in vitro* system during cell line development: growth factor independence, apoptotic evasion and limitless replicative potential. We then generated several genetically modified HMECs carrying different oncogenes (Figure 1a). The constitutively active mutant form of EGF receptor (EGFR L858R) and the oncogenic K-ras (KRas) were stably integrated into HMECs to confer growth factor-independent proliferation ^42^, and the resulting modified cell lines are referred to as HMEC-EGFR and HMEC-KRas, respectively. To promote apoptotic evasion ^43^, we introduced a dominant negative mutant form of p53 (p53DD) to HMEC-KRas and generated HMEC-p53DD-KRas. Moreover, we developed another cell line harboring Simian Virus 40 Early Region (SV40ER), which also attenuates apoptosis due to the inhibitory role of SV40 large T antigen (LT) on p53. We refer to the resulting cell line developed from HMEC-KRas as HMEC-SV40ER-KRas. The successful expression of these genetic elements was validated by Western blot (Figure S1). In addition, we also obtained an HMEC line carrying Simian Virus 40 large T antigen (SV40-LT), oncogenic H-ras (HRas) and human telomerase reverse transcriptase (hTERT) ^44^. In this cell line, HRas and SV40-LT are able to induce growth factor-independent proliferation and evasion of apoptosis ^45^, respectively. The introduction of hTERT further enhances cellular replicative potential, so the resulting cell line HMEC-hTERT-LT-HRas possesses all three characteristic properties of cancer ^1^ relevant to our *in vitro* system (Figure 1a).

**Figure 1.**
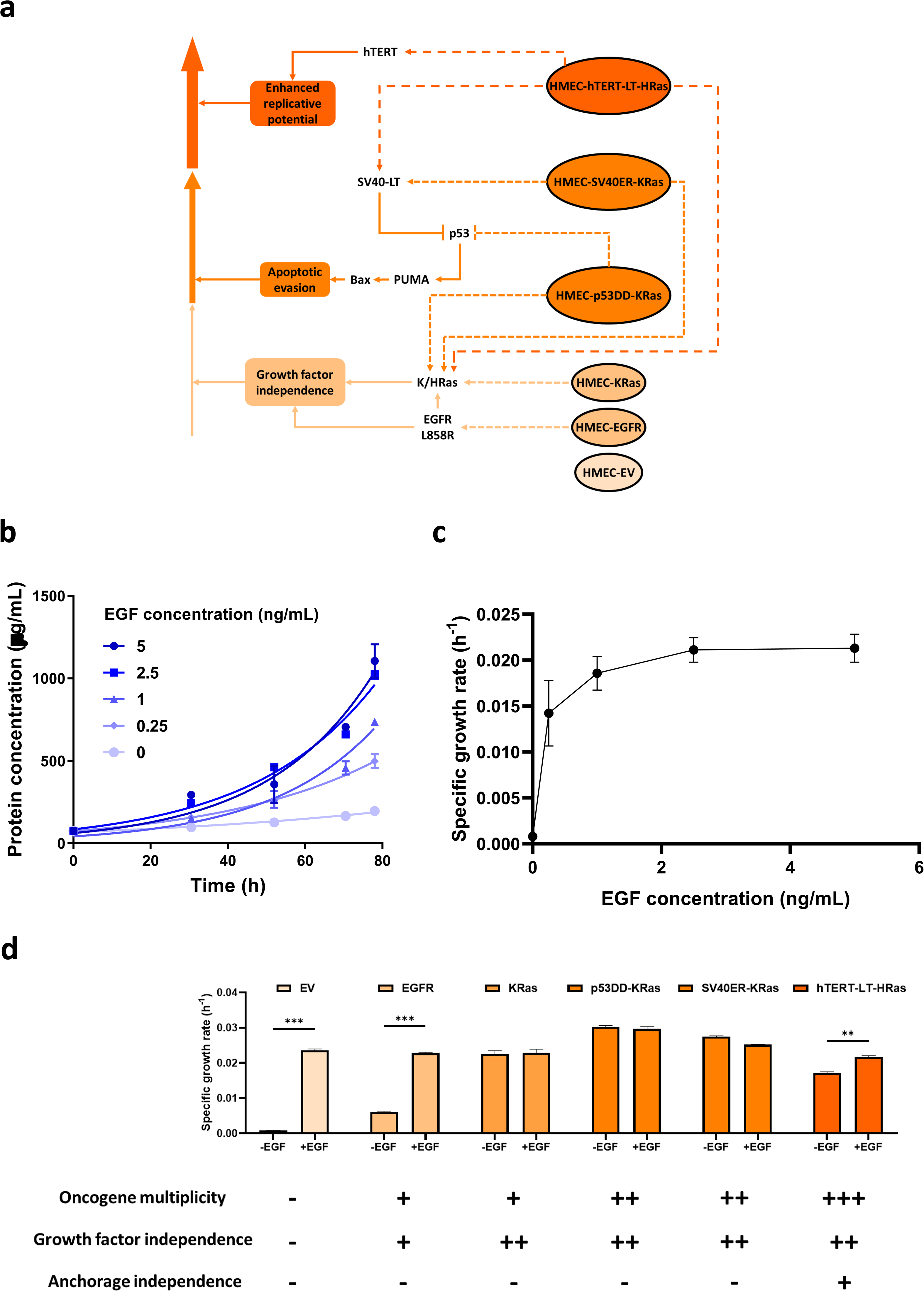
Development of a panel of HMECs with different combinations of oncogenes. **a**, Overexpression of defined oncogenes and dominant negative mutant forms of tumor suppressors in HMECs. Ovals indicate HEMC lines. Rounded rectangles indicate cancer-associated characteristics achieved during cell line development. Pointed and blunt-end arrows within the signaling pathway denote activation and inhibition, respectively. **b**, Proliferation of the normal HMECs in response to different EGF concentrations. **c**, Specific growth rate of the normal HMECs as a function of EGF concentration. **d**, Growth factor independence and other properties observed in genetically modified HMECs. Data are represented as mean ± SEM (n=3). **P<0.005. ***P<0.001.

Before characterizing the metabolic behavior of these cell lines, we first performed proliferation assays. Parental and empty vector-expressing HMECs exhibited exponential growth and were sensitive to EGF (Figure 1b). Based on exponential cell growth, we calculated specific growth rates at different EGF concentrations. EV cells proliferated faster at higher EGF concentrations, and the maximum specific growth rate was achieved at 5 ng/mL EGF (Figure 1c). We next performed proliferation assays in oncogene-containing HMECs with 5 ng/mL or no EGF (+/-EGF). Upon the integration of EGFR, cells began to acquire a small extent of growth factor independence, which means that they were able to actively proliferate in the absence of EGF. The uncontrolled proliferation was further enhanced in HMECs carrying KRas, p53DD-KRas, SV40ER-KRas and hTERT-LT-HRas. Accordingly, the panel of HMECs was ranked by increasing growth factor independence in Figure 1d. In addition, HMECs in this order also exhibited increasing oncogene multiplicity (including downregulation of tumor suppressor genes), with EV carrying no oncogene, EGFR and KRas harboring one, p53DD-KRas and SV40ER-KRas carrying two, and hTERT-LT-HRas possessing three oncogenic elements. Furthermore, previous work ^44^ has validated that HMEC-hTERT-LT-HRas is capable of undergoing anchorage-independent growth and forming tumors in immunodeficient mice, demonstrating the greatest level of tumorigenicity among all HMECs (Figure 1d). These HMECs with the same genetic background and defined oncogenic modifications enabled us to investigate whether cancer metabolism differs from normal proliferative metabolism in a well-controlled manner.

### Quantification of extracellular fluxes suggests dual control of metabolism by proliferation and oncogenotypes

We examined extracellular fluxes of glucose, lactate, glutamine and glutamate in our HMECs ^46^. The extracellular fluxes followed a similar trend as the proliferation rate: the difference between +/-EGF conditions was smaller in cell lines harboring more oncogenes (Figure 2a). We further calculated lactate yield on glucose and glutamate yield on glutamine by dividing the respective extracellular fluxes and accounting for the stoichiometry of converting glucose to lactate and glutamine to glutamate (Figure S2a). Interestingly, although the glutamate yield from glutamine exhibited a similar pattern as the extracellular fluxes, the lactate yield from glucose was not significantly different at varying oncogenotypes or in the presence or absence of EGF (Figure S2a). The only exception was the HMEC-EV -EGF condition, which showed a relatively small lactate yield. The invariant yield of lactate suggests the strong presence of the Warburg effect while cells are actively proliferating ^1, 27, 28^.

**Figure 2.**
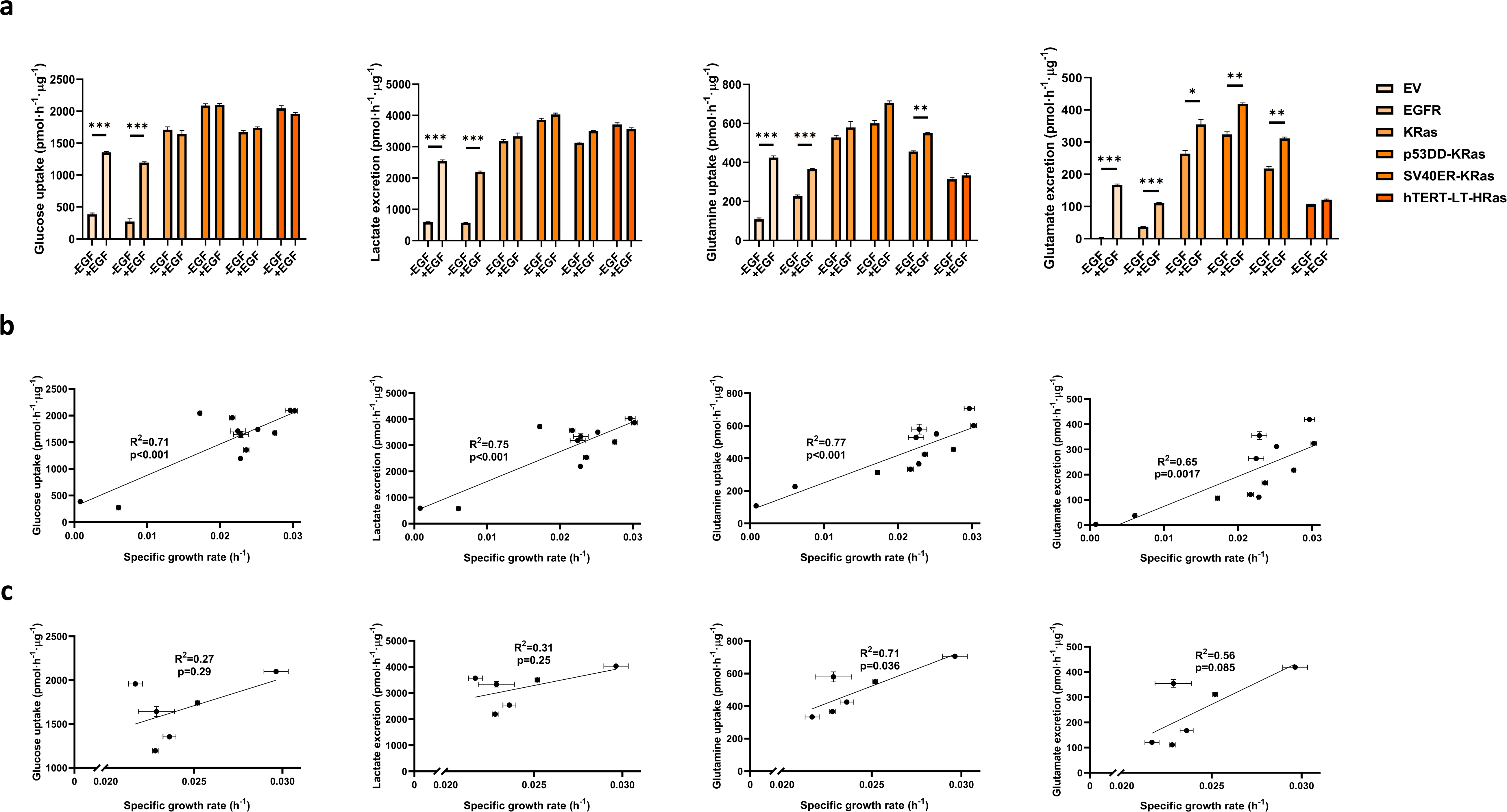
Extracellular fluxes were determined by both specific growth rates and oncogenotypes. **a,** Extracellular fluxes of glucose, lactate, glutamine and glutamate for HMECs at +/-EGF conditions. **b**, Extracellular fluxes of +/-EGF HMECs plotted against specific growth rates. **c**, Extracellular fluxes of only the +EGF HMECs plotted against specific growth rates. Data are represented as mean ± SEM (n=3). *P<0.05. **P<0.005. ***P<0.001.

Since the pattern of differential extracellular fluxes between +/-EGF conditions (Figure 2a) parallels that of growth (Figure 1 d), we sought to investigate whether growth rate regulates extracellular fluxes. To this end, we plotted extracellular fluxes against specific growth rates. All major extracellular fluxes (Figure 2b) and glutamate yield (Figure S2b) were higher at elevated growth, except for lactate yield, which stayed constant again presumably because of the Warburg effect (Figure S2b). Remarkably, the extracellular fluxes from both +/-EGF conditions tend to collapse onto a single regression line, which suggests the presence of direct control of extracellular fluxes by proliferation (Figure 2b). However, although the entire data set can be described by regression lines to some extent, a closer examination of the data points with comparable specific growth rates (0.02-0.025 h^-1^) suggests that extracellular fluxes are not solely determined by proliferation (Figure 2b). In fact, we constructed separate plots for HMECs grown at the +EGF condition (Figure 2c). The apparent proliferative control of extracellular fluxes, as indicated by the R^2^ and regression p-values, were reduced and increased respectively when considering only the +EGF condition (Figure 2c) compared to those in both +/-EGF conditions (Figure 2b and S3). Collectively, these data suggest that metabolism, at least manifested by extracellular fluxes, may be dually controlled by proliferation and specific oncogenotypes. In addition, proliferative control of metabolism was reduced in +EGF HMECs that shared similar growth rates.

### ^13^C-isotopic labeling reveals distinct substrate utilization patterns in the TCA cycle and *de novo* lipogenesis

To determine further metabolic changes, we performed ^13^C-isotopic labeling experiments by culturing HMECs in media separately containing three different tracers: uniformly ^13^C-labeled glucose (U-^13^C_6_-glucose); 1,2-^13^C-labeled glucose (1,2-^13^C_2_-glucose); and uniformly ^13^C-labeled glutamine (U-^13^C_5_-glutamine). Previous studies have shown that these three tracers give the best resolution of fluxes for glycolysis, PPP and the TCA cycle ^47^. Measurement of isotope enrichment of TCA cycle intermediates and amino acids revealed a distinct substrate utilization pattern in all HMECs used in our study. The canonical view of anaplerosis predicts strong enrichment of all TCA cycle intermediates by U-^13^C_6_-glucose. However, we observed a low level of enrichment for all U-^13^C_6_-glucose-derived metabolites of the TCA cycle. In fact, the majority of TCA cycle metabolites were weakly labeled by U-^13^C_6_-glucose, except for citrate, which was mainly labeled by two ^13^C atoms (M+2). This labeling pattern indicated that the ^13^C influx from glucose stopped at the point of citrate (Figure 3a). This conclusion was further corroborated by 1,2-^13^C_2_-glucose labeling, which generated similar profiles compared to those from U-^13^C_6_-glucose, consistent with the notion that M+2 citrate was the only metabolite moderately labeled by glucose tracers (Figure S4).

**Figure 3.**
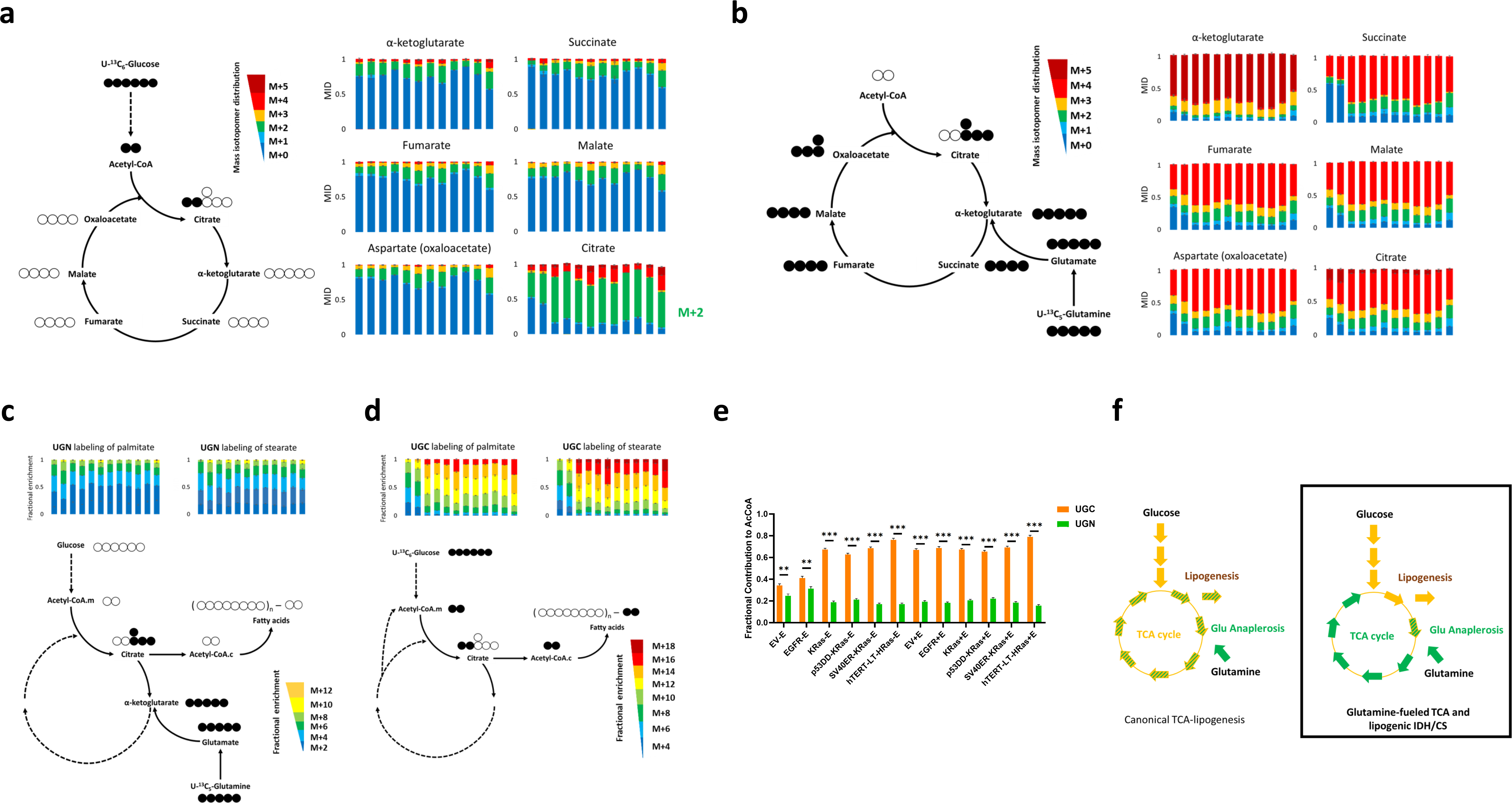
^13^C-isotopic labeling analysis revealed a truncated TCA cycle. **a**, U-^13^C_6_-glucose labeling pattern suggests weak incorporations of glucose-derived carbons into the TCA cycle except for citrate, the majority of which stays at M+2. **b**, U-^13^C_5_-glutamine labeling pattern suggests much stronger incorporations of glutamine-derived carbons into the TCA cycle. **c**, U-^13^C_5_-glutamine labeling pattern suggests weak incorporations of glutamine-derived carbons into *de novo* fatty acid synthesis. **d**, U-^13^C_6_-glucose labeling pattern suggests stronger incorporations of glucose-derived carbons into *de novo* fatty acid synthesis. **e**, Isotopomer spectral analysis (ISA) confirmed that glucose dominated lipogenesis over glutamine. **f**, ^13^C-isotopic labeling analysis revealed a distinct substrate utilization pattern within the TCA cycle and *de novo* lipogenesis in contrast to the canonical view. Columns along the axis of abscissas in each bar graph represent ^13^C-isotopic labeling data in the following order: EV -EGF, EGFR -EGF, KRas -EGF, p53DD-KRas -EGF, SV40ER-KRas -EGF, hTERT-LT-HRas -EGF; EV +EGF, EGFR +EGF, KRas +EGF, p53DD-KRas +EGF, SV40ER-KRas +EGF, hTERT-LT-HRas +EGF. Mass isotopomer distribution (MID) is shown in **a** and **b**. Fractional enrichment is shown in **c** and **d**. Open and filled circles in the schematics indicate ^12^C an ^13^C atoms within the metabolites, respectively. Acetyl-CoA.c and acetyl-CoA.m refer to cytosolic and mitochondrial acetyl-CoA, respectively. Data are represented as mean ± SEM (n=3). Abbreviations: UGC, U-^13^C_6_-glucose; UGN: U-^13^C_5_-glutamine; Glu, glutamate.

Intrigued by this stark distribution pattern of glucose-derived carbons, we further labeled cells with U-^13^C_5_-glutamine, hypothesizing that glutamine may be the primary carbon substrate fueling the TCA cycle. The crucial role of glutamine anaplerosis has been reported in various studies. In several types of cancer cells grown under hypoxia, glutamine provides carbon sources for *de novo* lipogenesis through the reversal of isocitrate dehydrogenase (IDH) ^36, 48, 49^. In contrast to reductive metabolism, in our HMECs, glutamine was primarily incorporated into the oxidative TCA cycle. Mass isotopomer distribution (MID) data showed that TCA intermediates were labeled much more strongly by U-^13^C_5_-glutamine than U-^13^C_6_-glucose. Across all cell lines, TCA cycle metabolites were approximately 50% enriched to the highest isotopomer states (M+4/5) when cultures were labeled by U-^13^C_5_-glutamine. Moreover, M+4 abundance was much greater than that of M+5 for citrate, indicating that the primary route of glutamine anaplerosis was oxidative cycling rather than reductive carboxylation ^50^. This data suggests that glutamine, but not glucose, is the primary carbon source that fuels the oxidative TCA cycle (Figure 3b).

In addition to glutamine anaplerosis, we also investigated the metabolic fate of glucose-derived carbons. Aside from citrate, TCA cycle metabolites were minimally labeled by U-^13^C_6_-glucose (Figure 3a). This labeling pattern suggests that glucose-derived fluxes are diverted away from the TCA cycle at the point of citrate. Because citrate initiates *de novo* lipogenesis, we analyzed the nonpolar metabolites from cells cultured in ^13^C-labeled substrates to identify the metabolic fate of citrate. The MIDs of palmitate and stearate suggest that glucose, rather than glutamine, was the main contributor to *de novo* fatty acid synthesis (Figure 3c-d). To further quantify the fractional contributions of glucose and glutamine to acetyl-CoA, a two-carbon unit recruited for *de novo* lipogenesis, we performed isotopomer spectral analysis (ISA) ^51^. ISA confirmed the lipogenic role of glucose relative to glutamine (Figure 3e and S5). The labeling and ISA results suggest that glucose-derived carbons were mostly utilized for *de novo* lipogenesis. Taken together, our ^13^C-isotopic labeling results identified a distinct substrate utilization pattern regarding the TCA cycle and *de novo* lipogenesis in HMECs. In all cell lines that we studied, the TCA cycle functions in a truncated fashion: glutamine supplies the majority of the carbon substrates for TCA cycle intermediates, while lipogenesis is maintained primarily by glucose rather than glutamine (Figure 3f).

### Quantification of intracellular metabolism via ^13^C-metabolic flux analysis (^13^C-MFA) corroborates dual control of metabolism by proliferation and oncogenotypes

In order to quantitatively investigate intracellular metabolism in HMECs, we performed ^13^C-metabolic flux analysis (^13^C-MFA). Briefly, ^13^C-MFA estimates metabolic fluxes by fitting them such as to most closely reproduce the measured metabolite isotopic enrichment patterns ^52^ (Figure 4a). The metabolic network model constructed for our ^13^C-MFA consists of major biochemical reactions within central carbon metabolism (Figure S6 and Table S1). ^13^C-MFA was performed under pseudo-steady state hypothesis, which was difficult to achieve in slowly growing cells. Accordingly, we report converged ^13^C-MFA flux results for all +EGF HMECs (Table S6-11) and those -EGF HMECs that still actively proliferated (KRas, p53DD-KRas, SV40ER-KRas and hTERT-LT-HRas, Table S2-5).

**Figure 4.**
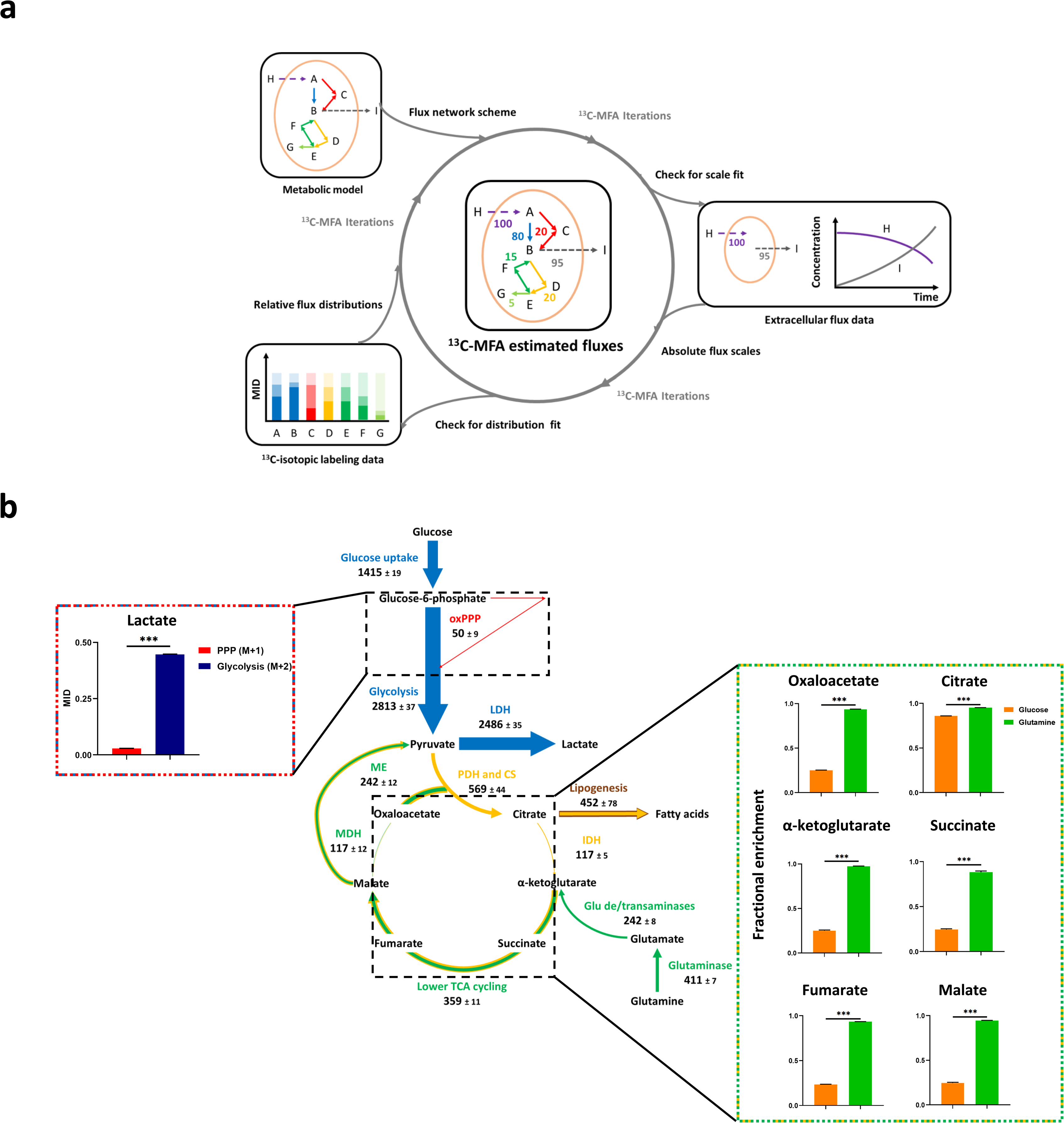
Introduction to ^13^C-MFA and consistency tests against the labeling data. **a**, Essential components and steps for ^13^C-metabolic flux analysis (^13^C-MFA). Based on a detailed metabolic network model, ^13^C-MFA determines absolute flux scales from extracellular flux data. ^13^C-isotopic labeling data is used to resolve intracellular flux distributions. The numerical algorithm works through iterations, during which ^13^C-MFA frequently checks the fit of experimentally determined extracellular fluxes and ^13^C-isotopic labeling data by values simulated by hypothesized flux distributions. The converged results are shown at the center of the schematic. **b**, ^13^C-MFA generated consistent results with the labeling data regarding the partition of fluxes between glycolysis and oxPPP and the anaplerotic contributions from glucose and glutamine. Flux results of +EGF HMEC-EV are shown here as an example. Unit of metabolic fluxes, pmol·h^-1^·μg^-1^. Uncertainties are shown as 95% confidence intervals. Data are represented as mean ± SEM (n=3). ***P<0.001. Abbreviations: oxPPP, oxidative pentose phosphate pathway; LDH, lactate dehydrogenase; ME: malic enzyme; MDH, malate dehydrogenase; PDH, pyruvate dehydrogenase; CS, citrate synthase; IDH, isocitrate dehydrogenase; TCA cycle, tricarboxylic acid cycle; Glu, glutamate.

We also performed several consistency tests to examine if model-estimated fluxes from the totality of labeling data were in agreement with local labeling results generated from specialized isotopic tracers. For example, culturing cells with 1,2-^13^C_2_-glucose allows estimation of relative fluxes between glycolysis and oxPPP. In brief, the glycolytic pathway retains both ^13^C atoms from 1,2-^13^C_2_-glucose and therefore generates M+2 labeled lactate as the main product. The oxidative branch of the PPP, on the other hand, involves a decarboxylation reaction which results in a loss of a ^13^C atom in the form of carbon dioxide, yielding M+1 labeled lactate ^53, 54^. Relative magnitude of glycolysis and oxPPP can therefore be deduced by comparing the MIDs of M+2 against M+1 lactate. This can be compared with the same ratio calculated from the MFA-estimated fluxes using different isotopic labels. In accordance with the ^13^C-labeling pattern indicating a significantly lower fraction of M+1 (0.03) as opposed to M+2 lactate (0.45), ^13^C-MFA results also suggested a minor diversion of fluxes into oxPPP (50 pmol·h^-1^·µg^-1^) from glycolysis (2813 pmol·h^-1^·µg^-1^). Moreover, we cultured HMECs with U-^13^C_5_-glutamine and U-^13^C_6_-glucose, and calculated fractional enrichments of major TCA intermediates. The labeling results indicated that the TCA cycle was primarily maintained by glutamine. Consistently, MFA also suggested stronger anaplerotic flux by glutamine than that by glucose (Figure 4b).

It has been hypothesized that metabolic behavior in cancer and normal proliferative cells can be regarded similarly ^1, 25, 26, 55^. To test this notion and investigate how proliferation affects metabolism, we plotted major intracellular fluxes against specific growth rates in all HMECs with converged results. The regression trend lines suggested that glycolysis and lactate excretion increased as cells grew faster, consistent with the Warburg Effect (Figure 5). As expected, the malic enzyme (ME) reaction was also enhanced, potentially regenerating NADPH for cellular redox needs ^56–58^ (Figure 5). Moreover, glutamine anaplerosis was enhanced when cells grew faster. This metabolic pattern further pinpoints the vital role of glutamine metabolism in maintaining the TCA cycle (Figure 3f). Additionally, ^13^C-MFA results suggested that PDH, rather than pyruvate carboxylase (PC), catalyzed the primary route of pyruvate entry into the TCA cycle (Figure S8), consistent with what has been reported in other cell lines ^2^. Importantly, although the overall trend lines suggested that metabolic fluxes were controlled by growth rate at some level, we also noticed non-negligible deviations of certain fluxes from the regression lines (Figure 5). Such deviations indicated that growth was not the only factor determining fluxes. In other words, metabolism is affected by both proliferation and oncogenotypes.

**Figure 5.**
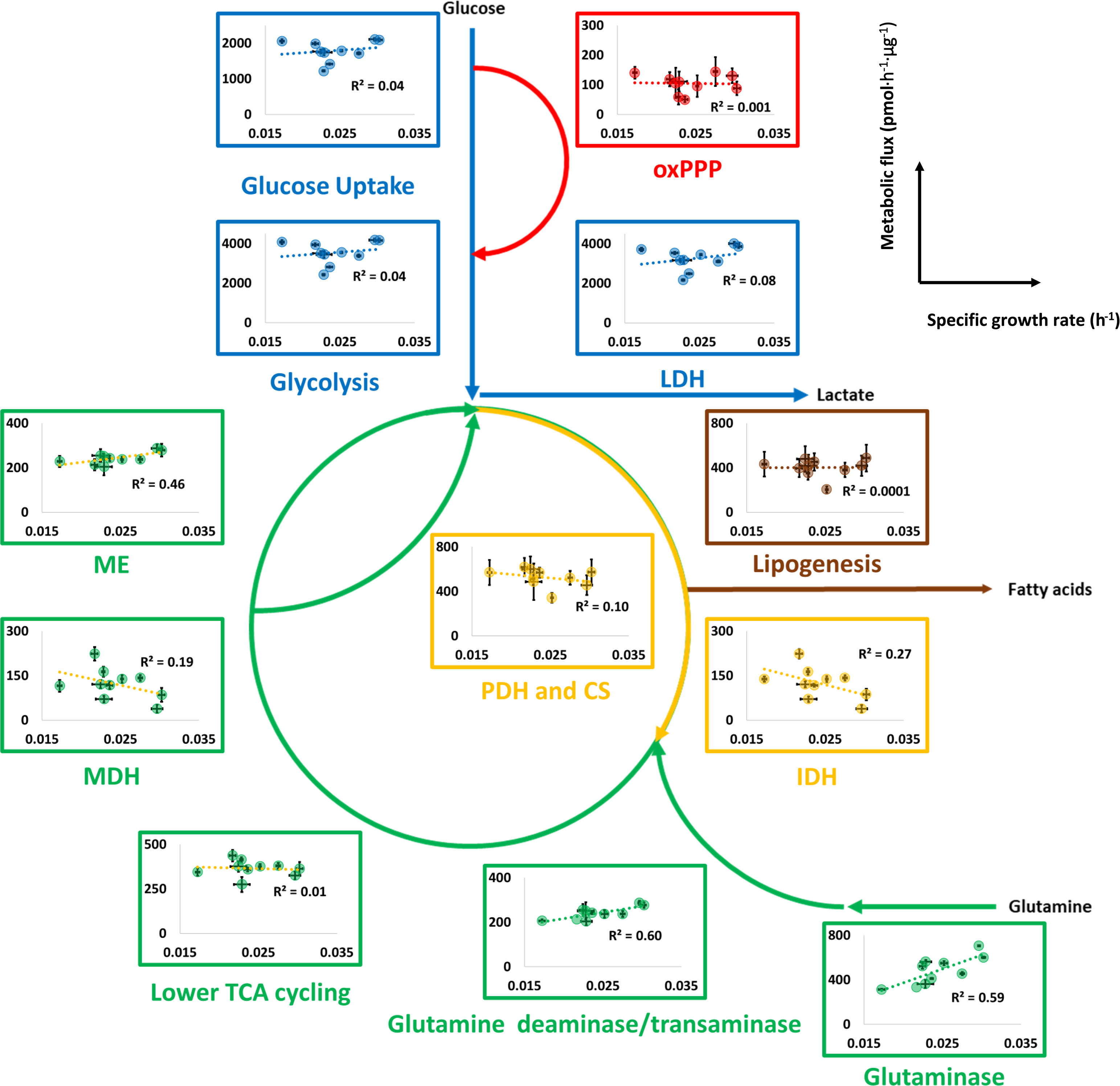
^13^C-MFA suggested that both intracellular and extracellular metabolisms may be dually controlled by proliferation and oncogenotypes. Data are represented as mean ± 95% confidence intervals. Abbreviations: oxPPP, oxidative pentose phosphate pathway; LDH, lactate dehydrogenase; ME: malic enzyme; MDH, malate dehydrogenase; PDH, pyruvate dehydrogenase; CS, citrate synthase; IDH, isocitrate dehydrogenase; TCA cycle, tricarboxylic acid cycle; Glu, glutamate.

### Normalizing fluxes against growth by introducing metabolic flux intensity (MFI)

Our ^13^C-MFA results (Figure 5) and extracellular fluxes (Figure 2) suggest that metabolism may be dually controlled by both proliferation and oncogenotypes. Indeed, there may be two modes of action through which metabolism is regulated: an indirect route through which oncogene-altered proliferation affects metabolism and a direct oncogenic effect independent of proliferative control (Figure S10).

The two modes of control were uncoupled by dividing metabolic fluxes by the specific growth rate (Figure S10). We term the resulting quantity as Metabolic Flux Intensity (MFI). When the MFI of a pathway increases under some conditions, this suggests that cells require a higher flux along the pathway to sustain the same level of growth, whereas lower MFI suggests that the corresponding metabolic pathway plays a less essential role in sustaining cellular growth. Therefore, the MFI of a pathway is an indicator of how strongly cells rely on the pathway to proliferate. By defining this new quantity, we can potentially assess the direct impact of oncogenotypes on metabolism independent of proliferation.

We thus calculated MFIs for all major metabolic pathways and constructed the plots of MFIs against specific growth rates (Figure 6). To isolate the impact of oncogenotype alone, we chose to focus on the +EGF HMECs. In addition, MFIs were averaged for oncogene-containing yet non-cancerous HMECs (EGFR, KRas, p53DD-KRas and SV40ER-KRas) to underscore the overall trend. We noticed that MFIs of certain metabolic pathways were increased in more cancerous lines, while other pathways remained unchanged (Figure 6). Interestingly, the glycolytic pathway and the lactate dehydrogenase (LDH) reaction exhibited enhanced MFIs, suggesting that the Warburg effect extends beyond the simple notion of proliferative upregulation of aerobic glycolysis (Figure 6). Instead, cells with higher oncogenic potential may need even greater glycolytic and LDH fluxes per growth rate to maintain proliferation. Importantly, pathways with the greatest MFIs were oxPPP, malate dehydrogenase (MDH) and isocitrate dehydrogenase (IDH). Based on the definition of MFI, these may be the most critical reactions for sustaining growth in HMEC-hTERT-LT-HRas relative to the control cell line (Figure 6).

**Figure 6.**
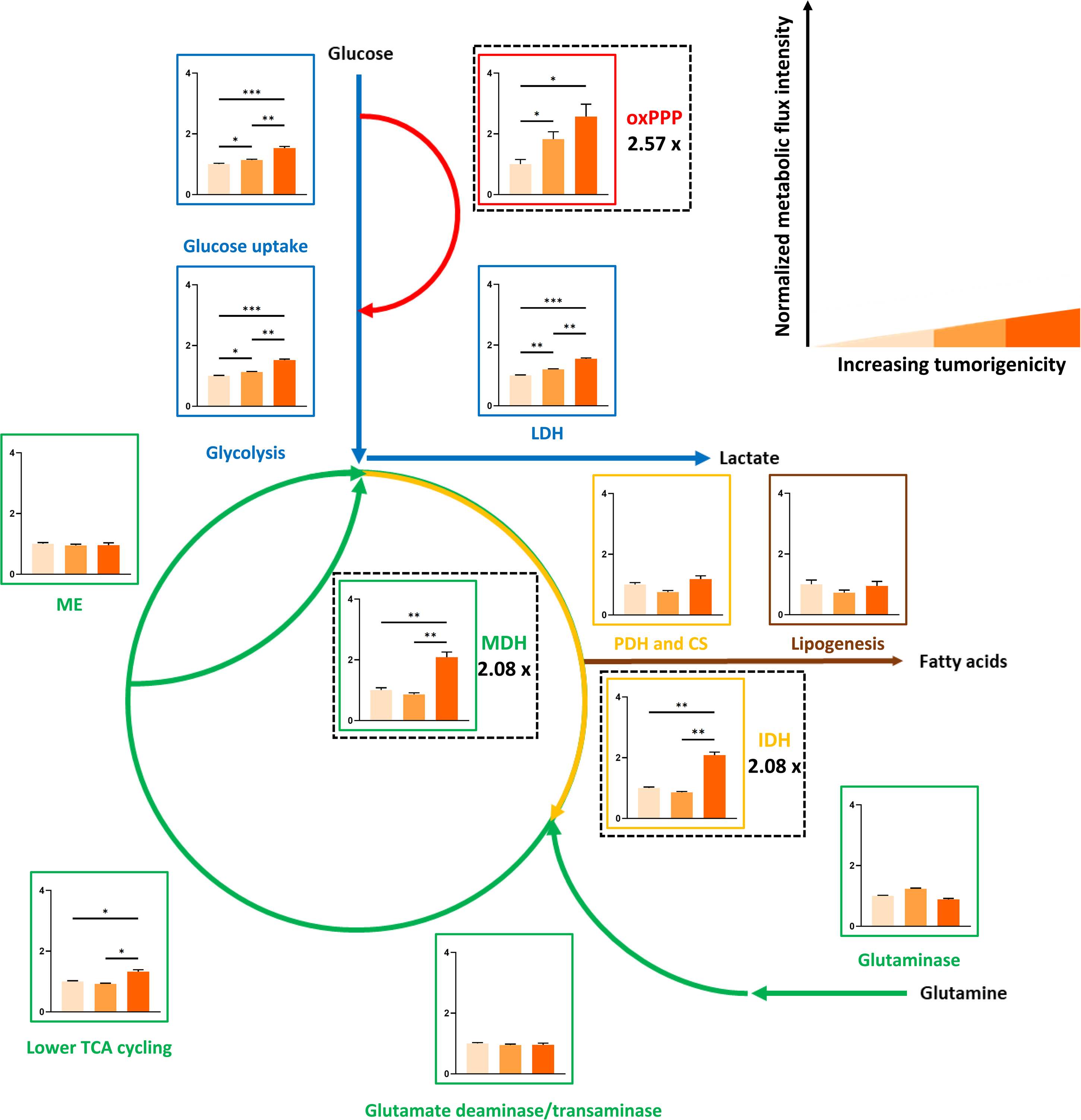
Quantitative ^13^C-MFI analysis validated that oncogenotypes directly impact metabolism independent of proliferative control. Normalized MFI results for all +EGF HMECs. Data are represented as mean ± 95% confidence intervals. *P<0.05. **P<0.005. ***P<0.001. Abbreviations: oxPPP, oxidative pentose phosphate pathway; MDH, malate dehydrogenase; IDH, isocitrate dehydrogenase; LDH, lactate dehydrogenase; ME: malic enzyme; PDH, pyruvate dehydrogenase; CS, citrate synthase; TCA cycle, tricarboxylic acid cycle.

### Assessing the therapeutic potential of targeting oxPPP, MDH and IDH

Our MFI analysis identified oxPPP, MDH and IDH as the most enhanced reactions in HMECs harboring hTERT-LT-HRas. Interestingly, these three pathways are essential sources regenerating NADH and NADPH. OxPPP is one of the primary routes of NADPH synthesis, which can also be replenished by ME (Figure 7a). However, ME exhibited statistically invariant MFIs across cell lines, whereas the reliance on oxPPP was higher in more cancerous HMECs (Figure 6). Additionally, MDH and IDH play an important role in maintaining the integrity of the TCA cycle, which functions as a central hub for supplying ATP, NADH and amino-anabolic precursors (Figure 7b). Therefore, inhibition of oxPPP, MDH and IDH may be selectively toxic in cancerous HMECs.

**Figure 7.**
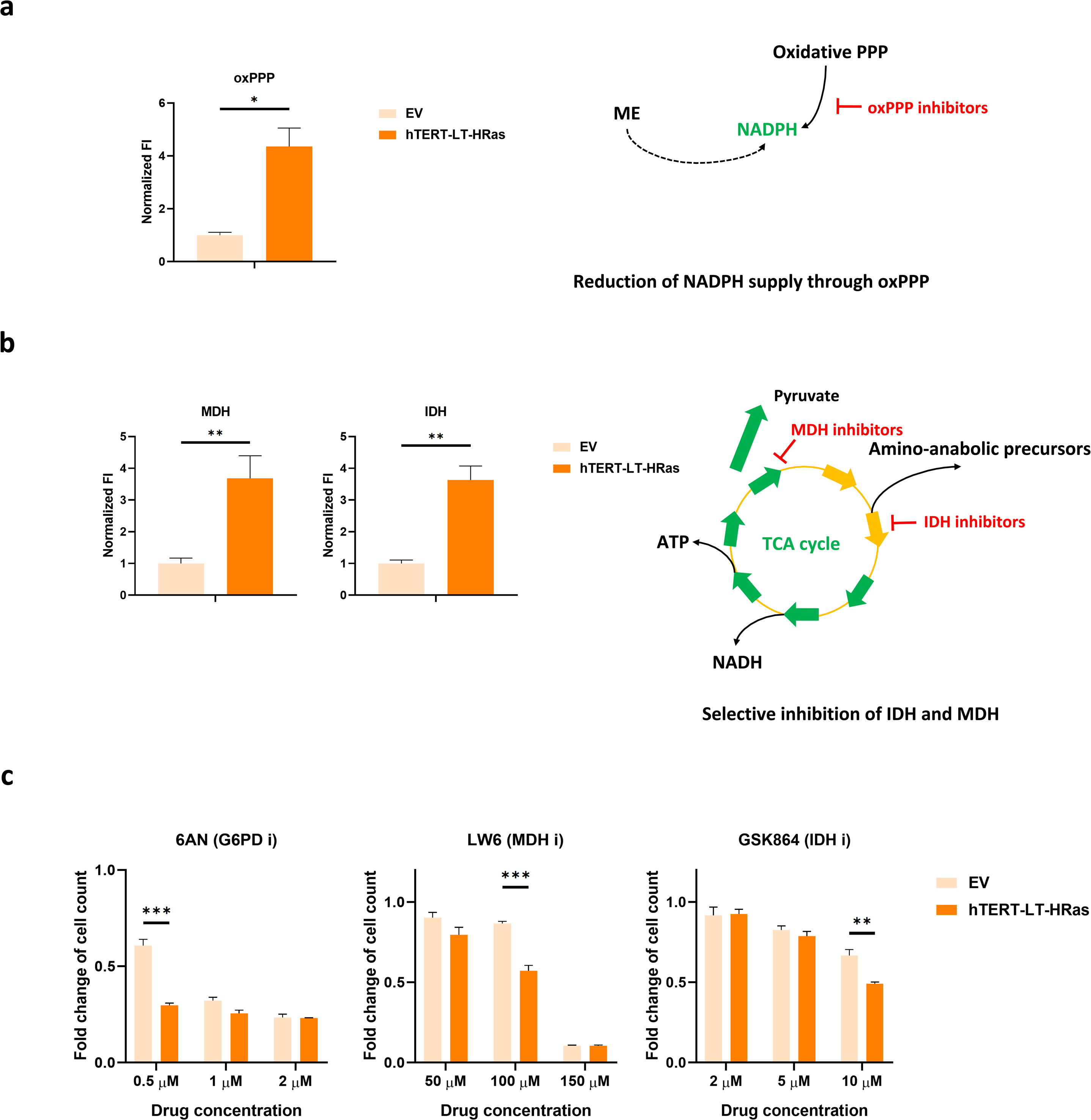
Therapeutic potentials of targeting oxPPP, MDH and IDH to selectively kill the tumorigenic HMEC as opposed to the normal proliferating counterpart. **a**, ^13^C-MFI analysis identified oxPPP as the most enhanced reaction in HMEC-hTERT-LT-HRas. Targeting oxPPP is expected to impair NADPH regeneration. **b**, ^13^C-MFI analysis identified MDH and IDH as one of the most enhanced reactions in HMEC-hTERT-LT-HRas. Targeting MDH and IDH is expected to impair NADH regeneration and the TCA cycle. **c**, Drug targeting results for oxPPP, MDH and IDH. Data are represented as mean ± 95% confidence intervals (**a**, **b**), and mean ± SEM (n=3) (**c**). *P<0.05. **P<0.005. ***P<0.001. Abbreviations: oxPPP, oxidative pentose phosphate pathway; ME: malic enzyme; MDH, malate dehydrogenase; IDH, isocitrate dehydrogenase; G6PD, glucose-6-phosphate dehydrogenase; 6AN, 6-aminonicotinamide.

To test these two strategies, we performed drug inhibition experiments for the tumorigenic HMEC-hTERT-LT-HRas against the control line HMEC-EV. We used 6-aminonicotinamide (6AN), LW6 and GSK864 to inhibit oxPPP ^59^, MDH ^60^ and IDH ^61^, respectively. The fold changes of cell count for drug-treated compared to DMSO-treated cells at different drug concentrations are shown in Figure 7c. We found that these inhibitors were selectively more toxic in the tumorigenic HMEC. Specifically, 0.5 μM LW6 or 10 μM GSK864 significantly reduced the proliferation of HMEC-hTERT-LT-HRas compared to the control cell line. It is worth noting that drug targeting of the metabolic pathways exhibiting statistically invariant MFIs might not be able to achieve selective toxicity (Figure S13). Therefore, pathways with the greatest MFIs in the tumorigenic line may serve as better drug targets. These experiments underscore the therapeutic potentials of targeting oxPPP, MDH and IDH for selectively killing cancerous HMECs.

## Discussion

We have demonstrated the use of ^13^C-isotopic labeling and MFA to quantitatively resolve the metabolic fluxes of HMECs with different combinations of oncogenes. Our data and analysis provide direct evidence that the metabolisms of cancer and normal proliferative cells differ and this can be the basis for identifying targets for selective cancer treatment. In addition, we found a distinct substrate utilization pattern involving a truncated TCA cycle and glucose-supported lipogenesis. Moreover, we introduced the concept of MFI to unveil proliferation-independent oncogenic metabolic rewiring that helped identify pathways like oxPPP, MDH and IDH as new drug targets with improved selective toxicity.

Our work further underscores the metabolic significance of glutamine anaplerosis. The fact that glutamine is metabolized as one of the major respiratory substrates in cancer cells has been previously reported ^62–64^. More recently, the important role of glutamine in supporting *de novo* lipogenesis has been elucidated ^36, 37, 49^. In contrast to these works, here we revealed another type of glutamine metabolism: a truncated TCA cycle for glutamine oxidation with glucose mainly supporting the lipogenic pathway. It is worth mentioning that a truncated TCA cycle has been reported in previous investigations. For example, glutamine can be utilized to fuel the TCA cycle in cancer cell lines harboring heterozygous IDH-1 mutations under hypoxia or mitochondrial inhibition ^65^. Moreover, cell lines exhibiting a truncated TCA cycle exhibited reduced proliferation due to their inability to perform a fully functional reductive glutamine metabolism along IDH ^65^. In contrast, our results show that the presence of a truncated TCA cycle does not undermine cellular growth in HMECs (Figure 1 b-d). Instead, further truncation was observed in the cell lines that proliferated faster (Figure 3e), suggesting that this utilization pattern may be regulated by proliferation. In line with our results, the truncated TCA cycle and glucose-supported lipogenesis have also been reported in glioblastomas, and such rewired metabolism does not impair cancer cell proliferation either ^2^. Importantly, our work shows that the truncated TCA cycle is present in all HMECs being studied, indicating that this metabolic pattern is not unique to cancer cells. Furthermore, we employed ^13^C-MFA to fully resolve all major metabolic pathways and quantified fluxes within the truncated TCA cycle and *de novo* lipogenesis. This quantitative depiction provides opportunities to further investigate this metabolic pattern in depth. For example, the mechanism by which this metabolic phenomenon occurs is still unclear at this point and remains to be elucidated by future work. These rewired energetics may expose potential metabolic vulnerabilities that can be exploited for therapeutic intervention to treat cancer. This idea is in line with the therapeutic efforts to target both glucose and glutamine metabolism for enhanced overall effectiveness of chemotherapy ^66–68^.

Our quantitative flux results enabled us to address the question of whether cancer and normal proliferative metabolisms are different. One of the hallmarks of cancer – the Warburg effect – has been proposed to be shared by both cancer and proliferating cells ^1, 27, 28^. However, there has been no study directly investigating whether and how cancer metabolism is different from proliferative metabolism beyond the Warburg effect. In fact, it would be difficult to eliminate confounding factors such as different cell types and genetic backgrounds if common existing cancer cell lines were used in such studies. These concerns prompted us to develop a new panel of HMECs that shared the same original genetic background by modifying defined genetic elements. More importantly, we believe that the main challenge to study the difference between cancer and normal proliferative metabolism is to decouple the growth-independent impact by oncogenotypes from that by altered growth phenotypes ^69^.

To decouple these two modes through which cancer metabolism is manifested, we introduced a new quantity called metabolic flux intensity (MFI) obtained by dividing metabolic fluxes by the specific growth rate. In this way, the effect of growth on metabolism is normalized when one compares MFIs across different cell lines. MFI essentially serves as an indicator gauging the importance of a certain metabolic pathway in sustaining cellular growth. MFIs are different across our HMEC variants, which suggests that proliferation is not the only factor governing metabolic behavior.

MFI analysis helps in selecting targets for treating cancer. Based on the definition of MFI, metabolic pathways with higher MFIs are relied upon by the cell more heavily than those with lower MFIs for supplying metabolites and energy for growth. Therefore, pathways with higher MFIs can be therapeutically more relevant in exerting selective toxicity on cancer cells. MFI analysis emphasizes the notion of selective toxicity because it compares flux changes independent of proliferative control of metabolism. Therefore, MFI may be more suitable for identifying potential drug targets.

Since oxPPP, MDH and IDH are responsible for producing NADPH and NADH ^70^, cofactor regeneration may be one of the key constraints limiting cancer cell growth. In line with this idea, inhibition of these reactions rendered a more drastic reduction of proliferation in the tumorigenic HMEC line. Previous studies also reported similar promising results ^71–74^. We believe that MFI analysis can be helpful in identifying potential drug targets in other types of cancer.

## Significance

In order to distinguish the metabolism of cancer and normal growing cells, we performed stable isotope tracing and metabolic flux analysis in a panel of human mammary epithelial cells (HMECs) harboring different oncogenes. We identified oxidative pentose phosphate pathway, malate dehydrogenase and isocitrate dehydrogenase as the most activated metabolic reactions and potential drug targets in the tumorigenic HMEC. Moreover, we introduced a new quantity termed metabolic flux intensity to describe metabolic rewiring independent of proliferative control. Our quantitative metabolic analysis provides direct evidence that metabolism is dually controlled by proliferation and oncogenes, thus supporting efforts to develop effective cancer treatment by selectively targeting metabolic pathways.

## Acknowledgements

We thank W. Hahn, O. Gjoerup, J. Deangelo and W. Kim for generously granting us one of the cell lines to study. We also thank J. Park, B. Woolston, B. Pereira, Z. Li, E. Lien, K. Abbott and A. Muir for discussions. We would like to further thank X. Lian and X. Bao for the suggestions on manuscript revision. The authors are supported by NIH R01 CA160458.

## Author contributions

Conceptualization, W.D., M.A.K. and G.S.; Methodology, W.D., M.A.K. and G.S.; Software, W.D., M.A.K. and N.L.; Validation, W.D., P.C. and J.L.C.; Formal analysis, W.D., M.A.K., S.J.M. and N.L.; Investigation, W.D. M.A.K. and S.J.M.; Resources, W.D., P.C., J.L.C., C.B. and G.S.; Writing – Original Draft, W.D. M.A.K. and P.C.; Writing – Review and Editing, W.D., M.A.K., S.J.M., P.C., J.L.C., N.L., J.K.K., M.G.V.H., O.I., H.D.S. and G.S., Supervision, J.K.K., M.G.V.H., O.I., H.D.S. and G.S.; Project Administration, G.S.; Funding Acquisition, G.S.

## Materials and methods

### Cell culture conditions

All cell lines were obtained from ATCC, tested for mycoplasma and cultured at 37 °C and 5% CO_2._ The MCDB 170 medium was used for regular subculture on 10-cm tissue-culture treated polystyrene dishes (Corning). The complete formulation of the MCDB 170 medium includes a mammary epithelial basal medium (MEBM) (Lonza) supplemented with 5 µg/mL insulin, 0.5 µg/ml hydrocortisone, 5 µg/mL transferrin, 0.07 mg/mL bovine pituitary extract (BPE) (Hammond), 10 uL isoproterenol and 5 ng/mL EGF (Peprotech). Trypsin-EDTA (0.25%) (Thermo Fisher) was used as the dissociation reagent to passage cells upon 70-80% confluence for up to 4 passages. Fresh media (15 mL per plate) was replaced every 2-3 days. Cells were plated and tested on 6-well plates (Corning) for the experiments determining growth rates and extracellular fluxes. Dulbecco’s Modified Eagle’s Medium (DMEM) (without glucose, glutamine and sodium pyruvate, Corning) was used as the basal medium (2.5 mL per well) supplemented with other MCDB 170 components, 10 mM glucose, 4 mM glutamine, 0.1 mM ethanolamine, 0.1 mM phosphoethanolamine and 10 mM HEPEs. All components were obtained from Sigma unless otherwise noted.

### Transfection/infection and drug selection for HMECs

HMEC 184A1 was used as the parental cell line to develop all HMEC variants, except for HMEC-hTERT-LT-HRas, which was kindly provided by Dr. W. Hahn (Dana-Farber Cancer Institute). An empty vector, K-ras G12V, EGFR L858R, p53DD and SV40ER (Addgene) were γ-retroviral infections under the CMV promoter. HEK 293 T cells at 40-60% confluence were transfected with 5.25 µg pBABE-puro EV, pBABE-puro K-ras G12V, pBABE-puro EGFR L858R, pBABE-neo p53DD or pBABE-neo SV40ER, 4.725 µg gag/pol vector, 0.525 µg VSV-G and 31.5 µL X-tremeGENE HP DNA Transfection Reagent (Roche). All components were premixed and equilibrated for 15 min at 25 °C before being transferred in a dropwise manner into HEK 293 T cells grown in a 10-cm dish with 10 mL DMEM and 10 vol % fetal bovine serum (FBS) (Sigma SAFC). Media containing γ-retroviruses was harvested after 36-48 h incubation at 37 °C and 5% CO_2_ and was passed through 0.45 µm filters (Pall). The retentate (∼10 mL) containing γ-retroviruses was then mixed with 8 μg/mL Polybrene (Millipore) and added into HMECs at 20-30% confluence grown in a 10-cm dish. After 24-36 hours, the spent media with retroviruses was aspirated and the plate was washed by 10 mL PBS (Corning) for three times.

HMEC-EV, HMEC-EGFR and HMEC-KRas with successful integrations of retroviral vectors were selected with 0.5 µg/mL puromycin. HMEC-p53DD-KRas and HMEC-SV40ER-KRas were selected with 0.5 µg/mL puromycin and subsequently with 400 µg/mL G418. The concentrations of these drugs were verified to be able to kill all mock-infected HMECs. To avoid bias due to position effect, the final population of each cell line was composed of at least 25 independent clones. HMEC lines after drug selection were then cryopreserved by mixing 1.5-3×10^6^ cells with 1mL freezing media containing 75 vol % MEBM, 15 vol % FBS and 10 vol % glycerol (Sigma).

### Western blots

Cells were lysed in RIPA buffer (Thermo) containing protease and phosphatase inhibitor cocktails (Bimake B14001 and B15001-B). Protein concentration was quantified by BCA assay (Thermo Fisher). Equal amounts of protein were run on 4-20% Tris-Glycine gels (Invitrogen) and transferred to PVDF membranes. Membranes were blocked with 5% bovine serum albumin and probed with the following primary antibodies: β-actin (Sigma, Cat. No. A1978), EGFR (Thermo Fisher, Cat. No. MS-400-P1), ERK1/2 (Cell Signaling, Cat. No. 9102) phospho-ERK1/2 (Cell Signaling, Cat. No. 4370), p53 (BD Biosciences, Cat. No. 554293), SV40 T Ag (Santa Cruz, Cat. No. SC-147), α-Tubulin (Cell Signaling, Cat. No. 3873). Blots were imaged using Luminata Western HRP substrate (EMD Millipore).

### Determinations of specific growth rates and extracellular fluxes

Protein concentrations for cells within 6-well plates were quantified by BCA assay (Thermo Pierce). Specific growth rates (µ) were determined by the following equation:

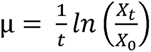

where *X_t_* and *X*_0_ are the protein concentrations at time 0 and time t, respectively. The initial cell seeding densities were 80×10^3^ and 35×10^3^ per well for -EGF and +EGF conditions, respectively. MCDB 170 medium was used during seeding (D-3), and DMEM with MCDB 170 components at +/-EGF conditions was used after 24 hours during media change (D-2), preceded by a two-time wash with PBS. Spent media was collected and new media was replaced after another 48 hours following a one-time wash with 2mL/well PBS (D0). Another 48 hours later, the spent media was harvested, and the cells (D2) were first washed once by 2 mL/well PBS. Then 500 µL/well of cold RIPA buffer (Thermo) was pipetted into each well, and the 6-well plates were shaken briefly and placed at 4 °C for overnight incubation. Cell lysates were then harvested, vortexed and centrifuged for the retrieval of supernatant. Protein concentration within each well was then measured in 96-well plates through colorimetric detection and quantification according to the standard protocol of BCA assay.

Concentrations of extracellular glucose, lactate, glutamine and glutamate in spent media were then measured by Yellow Spring Instruments (YSI) 2950. Based on the equation governing extracellular fluxes during the exponential growth phase, the extracellular fluxes for glucose, lactate and glutamate were determined according to the following equation ^46^:

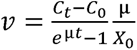

where *v* is the extracellular flux of a metabolite, *C_t_* the concentration of that metabolite at time t, *C*_0_ the concentration of that metabolite at time 0, μ the specific growth rate and *X*_0_ the initial protein content at time 0.

Due to the fact that glutamine is unstable and undergoes spontaneous degradation in normal cell culture condition, the extracellular fluxes for glutamine were determined based on the following equation with an additional term of decay constant k ^46^:

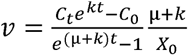

where the first-order decay constant k was determined based on the following equation:

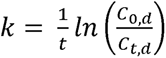

where *C_t,d_* and *C*_0*,d*_ are the glutamine concentrations in media without cells at time t and time 0, respectively.

### ^13^C-isotopic labeling experiments and intracellular metabolite extraction

^13^C-isotopic tracers (U-^13^C_6_-glucose, 1,2-^13^C_2_-glucose and U-^13^C_5_-glutamine, Cambridge Isotope Laboratories) were administered at the same concentrations as the unlabeled counterparts (10 mM for the glucose tracers and 4 mM for the glutamine tracer). The tracers were used one at a time at full enrichment. For example, 10 mM of U-^13^C_6_-glucose was used with 4 mM unlabeled glutamine in the U-^13^C_6_-glucose tracing experiment. The media formulation is exactly the same as that used in the growth and extracellular flux experiments. The experimental procedures are the same as that for the growth and extracellular flux experiments until D2. Instead of harvesting at D2, we prolonged the ^13^C-istopic labeling duration to 72 hours (D3) after the second media change (D0) to allow for more sufficient time to reach metabolic and isotopic steady states.

To harvest metabolites with minimal metabolic perturbations, we immediately placed the 6-well plates on ice once the plates were out of the incubator. Spent media was then quickly aspirated, and each well was washed once by cold (0-4 °C) saline, followed by addition of 500 µL freezing cold (−20 °C) methanol. Next, 300 µL cold (0-4 °C) milliQ water containing 2 µg norvaline (Sigma) as an internal standard was pipetted to each well. Cells were then quickly scrapped off from the wells by pipette tips in the presence of the liquid mixture, and the mixture was transferred to microcentrifuge tubes. These steps were performed one at a time for each well. Metabolite extraction was then performed by adding 600 µL freezing cold (−20 °C) chloroform to each microcentrifuge tube and then quickly vortexing the mixture at 4 °C for 10 min, followed by a centrifugation at 21000×g for 10 min. After these steps, the mixture was separated into two phases with the top and bottom layers containing polar and nonpolar metabolites, respectively. The two layers were then retrieved and placed into separate microcentrifuge tubes, dried by air and then stored at −80 °C for less than 1-2 weeks before being derivatized and loaded onto GC/MS.

### Metabolite derivatization and GC/MS analysis

Polar metabolites were derivatized by incubating dried samples in each microcentrifuge tube with 15 µL methoxyamine in pyridine (MOX) (Thermo) at 40 °C for 1.5 hours, followed by another incubation with 20 µL N-(tert-butyldimethylsilyl)-N-methyl-trifluoroacetamide with 1% tert-Butyldimethylchlorosilane (TBDMS) (Sigma) at 60 °C for 1 hour. The derivatized mixture was then briefly vortexed, centrifuged and transferred into polypropylene GC/MS vials (Agilent). Nonpolar metabolites were treated by incubating dried samples in each microcentrifuge tube with 500 µL methanol with 2 vol % sulfuric acid at 60 °C for 3 hours. Next, 600 µL hexane and 175 µL saturated sodium chloride solution were added, and the mixture was vortexed at room temperature for 30 min and subsequently centrifuged at 21000×g for 1 min. The resulting mixture was separated into two phases, and the top phase containing nonpolar metabolites was retrieved into a separate microcentrifuge tube, dried by air, reconstituted by 30-50 µL hexane and transferred into amber glass GC/MS vials with glass inserts (Agilent).

Agilent 6890N GC and 5975B Inert XL MS were used for polar metabolite analysis. The column for the 6890N GC is Agilent J&W DB-35ms (35%-phenyl-methylpolysiloxane, mid-polarity) and the electron ionization mode with 70 eV was used for the 5975B MS. Chromatography grade helium (Airgas) at 1mL/min flow rate was used as the carrier gas. The inlet temperature for the 6890N GC was set to 270 °C, and the oven temperature was first maintained at 100 °C, and ramped to 300 °C at a speed of 2.5 °C/min. Samples of either 1 or 2 µL were injected into the instrument with either split or splitless mode based on sample abundances. The scan mode with a detection range of 150-625 m/z was used for all measurements. Mass isotopomer distributions (MIDs) have been corrected for natural abundance.

Agilent 7890B GC and 5977B MS were used for nonpolar metabolite analysis. The column for the 7890B GC is Agilent J&W HP-5ms (5%-phenyl-methylpolysiloxane, nonpolar) and the electron ionization mode with 70 eV was used for the 5977B MS. Ultra high purity grade helium (Airgas) at 3 mL/min was used as the carrier gas. The inlet temperature for the 7890B GC was set to 280 °C, and the oven temperature was first maintained at 165 °C and then ramped to 226 °C at a speed of 2 °C/min. Samples of 1 µL were injected into the instrument with the splitless mode. The scan mode with a detection range of 200-400 m/z was used for all measurements. MIDs have been corrected for natural abundance.

### ^13^C-metabolic flux analysis (^13^C-MFA) and isotopomer spectral analysis (ISA)

An elementary metabolite unit (EMU)-based software Metran coded within MATLAB (MathWorks) was used to perform ^13^C-MFA ^52, 75^ and ISA ^36, 51^. Experimentally determined extracellular fluxes of glucose, lactate, glutamine and glutamate, as well as the ^13^C-labeling data were used in conjunction with a metabolic reaction model (Figure S3) to generate ^13^C-MFA results. The 95% confidence intervals for metabolic fluxes were obtained by performing parameter continuation on converged flux results ^76^. The critical steps of ISA are explained in Figure S5.

### Inhibition of oxPPP, MDH and IDH

HMEC-hTERT-LT-HRas and HMEC-EV were seeded at 35×10^3^ per well with 5ng/mL EGF in 6-well plates. The small molecule drugs 6-aminonicotinamide (6AN), LW6 and GSK864 at different concentrations were administered in the usual MEBM-based MCDB 170 medium.

DMSO was used to prepare the initial concentrated drug solutions. Cell counts were measured by Cellometer (Nexcelom) 48 hours after seeding.

### Definition of quantities

Fractional enrichments (also known as molar percent enrichment ^77, 78^) from ^13^C-isotopic tracers were calculated using MID data corrected for natural abundance based on the following equation:

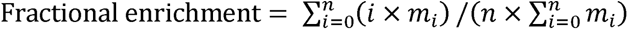

where *n* is the number of carbon atoms in a metabolite, *m_i_* the abundance of a mass isotopomer and *i* the labeling state (M+i) of a mass isotopomer.

Metabolic flux intensities (MFIs) were defined and calculated based on the following equation:

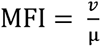

where *v* and μ refer to metabolic flux (pmol·h^-1^·µg^-1^) and specific growth rate (h^-1^), respectively.The unit of MFI is therefore pmol·μg^-1^.

## References

1. Hanahan, D. & Weinberg, R. A. Hallmarks of cancer: The next generation. Cell 144, 646– 674 (2011).

2. DeBerardinis, R. J. et al. Beyond aerobic glycolysis: transformed cells can engage in glutamine metabolism that exceeds the requirement for protein and nucleotide synthesis. Proc. Natl. Acad. Sci. U. S. A. 104, 19345–19350 (2007).

3. Hsu, P. P. & Sabatini, D. M. Cancer cell metabolism: Warburg and beyond. Cell 134, 703–707 (2008).

4. Liberti, M. V. & Locasale, J. W. The Warburg Effect: How Does it Benefit Cancer Cells? Trends Biochem. Sci. 41, 211–218 (2016).

5. Bensinger, S. J. & Christofk, H. R. New aspects of the Warburg effect in cancer cell biology. Semin. Cell Dev. Biol. 23, 352–361 (2012).

6. Koppenol, W. H., Bounds, P. L. & Dang, C. V. Otto Warburg’s contributions to current concepts of cancer metabolism. Nat. Rev. Cancer 11, 325–337 (2011).

7. Tran, T. Q. et al. α Ketoglutarate attenuates Wnt signaling and drives differentiation in colorectal cancer. *Nat*. Cancer 1, 345–358 (2020).

8. Keibler, M. A. et al. Differential substrate use in EGF- and oncogenic KRAS-stimulated human mammary epithelial cells. FEBS J. 1–21 (2021).

9. Warburg, O. On the origin of cancer cells. Science (80-.). 123, 309–314 (1956).

10. Vander Heiden, M. G., et al. Metabolic Pathway Alterations that Support Cell Proliferation. Cold Spring Harb. Symp. Quant. Biol. 76, 325–334 (2011).

11. Pacold, M. E. et al. A PHGDH inhibitor reveals coordination of serine synthesis and one-carbon unit fate. Nat. Chem. Biol. 12, 452–458 (2016).

12. Diaz-Moralli, S. et al. A key role for transketolase-like 1 in tumor metabolic reprogramming. Oncotarget 7, 51875–51897 (2016).

13. Ahn, W. S. et al. Glyceraldehyde 3-phosphate dehydrogenase modulates nonoxidative pentose phosphate pathway to provide anabolic precursors in hypoxic tumor cells. AIChE J. 64, 4289–4296 (2018).

14. Man, C. H. et al. Proton export alkalinizes intracellular pH and reprograms carbon metabolism to drive normal and malignant cell growth. Blood 139, 502–522 (2022).

15. Dong, W., Rawat, E. S., Stephanopoulos, G. & Abu-Remaileh, M. Isotope tracing in health and disease. Curr. Opin. Biotechnol. 76, 102739 (2022).

16. Gaglio, D. et al. Oncogenic K-Ras decouples glucose and glutamine metabolism to support cancer cell growth. Mol. Syst. Biol. 7, 1–15 (2011).

17. Son, J. et al. Glutamine supports pancreatic cancer growth through a KRAS-regulated metabolic pathway. Nature 496, 101–105 (2013).

18. Dang, C. V, Kim, J., Gao, P. & Yustein, J. The interplay between MYC and HIF in cancer. Nat. Rev. Cancer 8, 51–56 (2008).

19. Liu, W. et al. Reprogramming of proline and glutamine metabolism contributes to the proliferative and metabolic responses regulated by oncogenic transcription factor c-MYC. Proc. Natl. Acad. Sci. 109, 8983–8988 (2012).

20. Forcina, G. C. et al. Ferroptosis Regulation by the NGLY1/NFE2L1 Pathway. Proc. Natl. Acad. Sci. U. S. A. 119, 1–11 (2022).

21. Isidor, M. S. et al. Insulin resistance rewires the metabolic gene program and glucose utilization in human white adipocytes. Int. J. Obes. 1–9 (2021) doi:10.1038/s41366-021-01021-y.

22. Vander Heiden, M. G. Targeting cancer metabolism: A therapeutic window opens. Nat. Rev. Drug Discov. 10, 671–684 (2011).

23. Ngoi, N. Y. L. et al. Targeting Cell Metabolism as Cancer Therapy. Antioxidants Redox Signal. 32, 285–305 (2020).

24. Méndez-lucas, A. et al. Identifying strategies to target the metabolic flexibility of tumours. Nat. Metab. 2, 335–350 (2020).

25. Ward, P. S. & Thompson, C. B. Metabolic Reprogramming: A Cancer Hallmark Even Warburg Did Not Anticipate. Cancer Cell 21, 297–308 (2012).

26. Schulze, A. & Harris, A. L. How cancer metabolism is tuned for proliferation and vulnerable to disruption. Nature 491, 364–73 (2012).

27. Vander Heiden, M. G., et al. Understanding the Warburg Effect : Cell Proliferation. Science (80-.). 324, 1029–1034 (2009).

28. Vander Heiden, M. G. & DeBerardinis, R. J. Understanding the Intersections between Metabolism and Cancer Biology. Cell 168, 657–669 (2017).

29. Ananieva, E. Targeting amino acid metabolism in cancer growth and anti-tumor immune response. World J. Biol. Chem. 6, 281–289 (2015).

30. Chen, Z., Liu, M., Li, L. & Chen, L. Involvement of the Warburg effect in non-tumor diseases processes. J. Cell. Physiol. 233, 2839–2849 (2017).

31. Vacanti, N. M. & Metallo, C. M. Exploring metabolic pathways that contribute to the stem cell phenotype. Biochim. Biophys. Acta 1830, 2361–2369 (2013).

32. Folmes, C. D. L., Dzeja, P. P., Nelson, T. J. & Terzic, A. Metabolic plasticity in stem cell homeostasis and differentiation. Cell Stem Cell 11, 596–606 (2012).

33. Agathocleous, M. & Harris, W. A. Metabolism in physiological cell proliferation and differentiation. Trends Cell Biol. 23, 484–492 (2013).

34. Christofk, H. R. et al. The M2 splice isoform of pyruvate kinase is important for cancer metabolism and tumour growth. Nature 452, 230–233 (2008).

35. Kim, B., Li, J., Jang, C. & Arany, Z. Glutamine fuels proliferation but not migration of endothelial cells. EMBO J. 36, 2321–2333 (2017).

36. Metallo, C. M. et al. Reductive glutamine metabolism by IDH1 mediates lipogenesis under hypoxia. Nature 481, 380–384 (2012).

37. Mullen, A. R. et al. Reductive carboxylation supports growth in tumour cells with defective mitochondria. Nature 481, 385–388 (2012).

38. Beroukhim, R. et al. The landscape of somatic copy-number alteration across human cancers. Nature 463, 899–905 (2010).

39. Locasale, J. W. et al. Phosphoglycerate dehydrogenase diverts glycolytic flux and contributes to oncogenesis. Nat. Genet. 43, 869–74 (2011).

40. Olivares, O. et al. Collagen-derived proline promotes pancreatic ductal adenocarcinoma cell survival under nutrient limited conditions. Nat. Commun. 8, 1–14 (2017).

41. Hanahan, D. & Weinberg, R. A. The hallmarks of cancer. Cell 100, 57–70 (2000).

42. Schlaeth, M. et al. Fc-engineered EGF-R antibodies mediate improved antibody-dependent cellular cytotoxicity (ADCC) against KRAS-mutated tumor cells. Cancer Sci. 101, 1080–1088 (2010).

43. Polyak, K., Xia, Y., Zweier, J. L., Kinzler, K. W. & Vogelstein, B. A model for p53-induced apoptosis. Nature 389, 300–305 (1997).

44. Elenbaas, B. et al. Human breast cancer cells generated by oncogenic transformation of primary mammary epithelial cells. Genes Dev. 15, 50–65 (2001).

45. Shen, Y. & Shenk, T. E. Viruses and apoptosis. Curr. Opin. Genet. Dev. 5, 105–111 (1998).

46. Murphy, T. A. & Young, J. D. ETA: Robust software for determination of cell specific rates from extracellular time courses. Biotechnol. Bioeng. 110, 1748–1758 (2013).

47. Metallo, C. M., Walther, J. L. & Stephanopoulos, G. Evaluation of 13C isotopic tracers for metabolic flux analysis in mammalian cells. J. Biotechnol. 144, 167–174 (2009).

48. Yoo, H., Antoniewicz, M. R., Stephanopoulos, G. & Kelleher, J. K. Quantifying reductive carboxylation flux of glutamine to lipid in a brown adipocyte cell line. J. Biol. Chem. 283, 20621–20627 (2008).

49. Gameiro, P. A. et al. In vivo HIF-mediated reductive carboxylation is regulated by citrate levels and sensitizes VHL-deficient cells to glutamine deprivation. Cell Metab. 17, 372– 385 (2013).

50. Dong, W., Keibler, M. A. & Stephanopoulos, G. Review of metabolic pathways activated in cancer cells as determined through isotopic labeling and network analysis. Metab. Eng. 43, 113–124 (2017).

51. Kelleher, J. K. & Nickol, G. B. Isotopomer Spectral Analysis: Utilizing Nonlinear Models in Isotopic Flux Studies. Methods Enzymol. 561, 303–330 (2015).

52. Antoniewicz, M. R., Kelleher, J. K. & Stephanopoulos, G. Elementary metabolite units (EMU): A novel framework for modeling isotopic distributions. Metab. Eng. 9, 68–86 (2007).

53. Dong, W., Moon, S. J., Kelleher, J. K. & Stephanopoulos, G. Dissecting Mammalian Cell Metabolism through 13C-And 2H-Isotope Tracing: Interpretations at the Molecular and Systems Levels. Ind. Eng. Chem. Res. 59, 2593–2610 (2020).

54. Carpenter, K. L. H. et al. 13C-labelled microdialysis studies of cerebral metabolism in TBI patients. Eur. J. Pharm. Sci. 57, 87–97 (2014).

55. Sun, W. et al. TKTL1 is activated by promoter hypomethylation and contributes to head and neck squamous cell carcinoma carcinogenesis through increased aerobic glycolysis and HIF1α stabilization. Clin. Cancer Res. 16, 857–866 (2010).

56. Fan, J. et al. Quantitative flux analysis reveals folate-dependent NADPH production. Nature 10, 298–302 (2014).

57. Jiang, P., Du, W., Mancuso, A., Wellen, K. E. & Yang, X. Reciprocal regulation of p53 and malic enzymes modulates metabolism and senescence. Nature 493, 689–693 (2013).

58. DeBerardinis, R. J. & Chandel, N. S. Fundamentals of cancer metabolism. Sci. Adv. 2, 1– 18 (2016).

59. Tsouko, E. et al. Regulation of the pentose phosphate pathway by an androgen receptor-mTOR-mediated mechanism and its role in prostate cancer cell growth. Oncogenesis 3, 1– 10 (2014).

60. Zhang, X. et al. Metformin and LW6 impairs pancreatic cancer cells and reduces nuclear localization of YAP1. J. Cancer 11, 479–487 (2020).

61. Calvert, A. E. et al. Cancer-Associated IDH1 Promotes Growth and Resistance to Targeted Therapies in the Absence of Mutation. Cell Rep. 19, 1858–1873 (2017).

62. Moreadith, R. W. & Lehninger, A. L. Purification, kinetic behavior, and regulation of NAD(P)+ malic enzyme of tumor mitochondria. J. Biol. Chem. 259, 6222–6227 (1984).

63. Parlo, R. A. & Coleman, P. S. Enhanced rate of citrate export from cholesterol-rich hepatoma mitochondria. The truncated Krebs cycle and other metabolic ramifications of mitochondrial membrane cholesterol. J. Biol. Chem. 259, 9997–10003 (1984).

64. Piva, T. J. & McEvoy-Bowe, E. Oxidation of glutamine in hela cells: Role and control of truncated TCA cycles in tumour mitochondria. J. Cell. Biochem. 68, 213–225 (1998).

65. Grassian, A. R. et al. IDH1 mutations alter citric acid cycle metabolism and increase dependence on oxidative mitochondrial metabolism. Cancer Res. 74, 3317–3331 (2014).

66. Wise, D. R. & Thompson, C. B. Glutamine Addiction: A New Therapeutic Target in Cancer. Trends Biochem Sci. 35, 427–433 (2011).

67. Tennant, D. A., Durán, R. V. & Gottlieb, E. Targeting metabolic transformation for cancer therapy. Nat. Rev. Cancer 10, 267–277 (2010).

68. Jin, L., Alesi, G. N. & Kang, S. Glutaminolysis as a target for cancer therapy. Oncogene 35, 3619–3625 (2016).

69. Fritz, V. & Fajas, L. Metabolism and proliferation share common regulatory pathways in cancer cells. Oncogene 29, 4369–4377 (2010).

70. Moon, S. J., Dong, W., Stephanopoulos, G. N. & Sikes, H. D. Oxidative pentose phosphate pathway and glucose anaplerosis support maintenance of mitochondrial NADPH pool under mitochondrial oxidative stress. Bioeng. Transl. Med. 5, 1–18 (2020).

71. Fujii, T., Khawaja, M. R., DiNardo, C. D., Atkins, J. T. & Janku, F. Targeting Isocitrate Dehydrogenase (IDH) in cancer. Discov. Med. 21, 373–380 (2016).

72. Levis, M. Targeting IDH: The next big thing in AML. Blood 122, 2770–2771 (2013).

73. Lee, K., et al. Identification of malate dehydrogenase 2 as a target protein of the HIF-1 inhibitor LW6 using chemical probes. Angew. Chem. Int. Ed. Engl. 52, 10286–10289 (2013).

74. Ramos-Montoya, A. et al. Pentose phosphate cycle oxidative and nonoxidative balance: A new vulnerable target for overcoming drug resistance in cancer. Int. J. Cancer 119, 2733– 2741 (2006).

75. Young, J. D., Walther, J. L., Antoniewicz, M. R., Yoo, H. & Stephanopoulos, G. An elementary metabolite unit (EMU) based method of isotopically nonstationary flux analysis. Biotechnol. Bioeng. 99, 686–99 (2008).

76. Antoniewicz, M. R., Kelleher, J. K. & Stephanopoulos, G. Determination of confidence intervals of metabolic fluxes estimated from stable isotope measurements. Metab. Eng. 8, 324–337 (2006).

77. Laqtom, N. N. et al. CLN3 is required for the clearance of glycerophosphodiesters from lysosomes. Nature 609, 1005–1011 (2022).

78. Wortham, M., et al. Integrated In Vivo Quantitative Proteomics and Nutrient Tracing Reveals Age-Related Metabolic Rewiring of Pancreatic β Cell Function. Cell Rep. 25, 2904–2918.e8 (2018).

